# Effects of chronic mild stress and CB1 receptor activation on hippocampal-dependent fear conditioning in female adolescent rats

**DOI:** 10.64898/2026.02.13.705785

**Authors:** Christian G. Reich, Angelica Ferraro, Philip Wig, Nicole Amada, Michael S. Weiss

## Abstract

Sex differences in responses to chronic stress are implicated in the higher prevalence of major depression and PTSD in females. Evidence of sex differences in endocannabinoid (eCB) physiology suggests that eCB signaling contributes to sexual disparities in fear conditioning and extinction. In adolescent male Sprague-Dawley rats, exposure to chronic-mild-unpredictable stress (CMS) resulted in enhanced trace-fear conditioning that was reversed by CB1 activation (Reich et al, 2013). In the present study, we assessed the effects of CMS and CB1 activation on hippocampal-dependent trace and contextual fear conditioning in adolescent female Sprague-Dawley rats. CMS exposure enhanced trace freezing behavior during memory recall compared to non-stress controls. This effect was not observed in contextually conditioned females. The CB1 receptor agonist, ACEA (0.1 mg/kg), administered prior to trace memory recall, but not prior to acquisition, significantly decreased freezing in both stress and non-stress females. ACEA significantly reduced baseline freezing behavior during trace memory recall in both stress and non-stress rats, however ACEA either 1) did not affect or 2) impaired short and long-term extinction in stress and non-stress females. In contextually conditioned females, ACEA decreased freezing during memory recall, although the effect was more robust in stress rats. ACEA impaired long-term contextual extinction in stress females while facilitating this in non-stress controls. However, ACEA had no effect or impaired short-term contextual extinction in both stress and non-stress groups. The results demonstrate that CMS enhances hippocampal-dependent episodic fear memories but has limited effects on contextual fear conditioning in female rats. These findings have implications in the use of medical cannabinoid treatment of disorders such as PTSD, as well as recreational cannabis use in adolescent/young adult females.

## Introduction

Psychological sex differences in cognitive and emotional behaviors are well documented over the past several decades. These differences may underlie the observation that women are more vulnerable to stress-related mental disorders than men (Jovanovic and Ressler, 2010; Roy-Byrne et al.). Epidemiological studies in humans and experimental studies in rodents suggest that sex differences in the hypothalamic-pituitary-adrenal (HPA) axis stress response are responsible for these disparities (Jovanovic and Ressler, 2010; Roy-Byrne et al.; Schroeder et al., 2018; Tiwari and Gonzalez, 2018). Exposure therapies (i.e. systematic desensitization and flooding) are ubiquitous treatments of Post-traumatic-stress disorder (PTSD), phobias and anxiety disorders (Rau et al., 2005; Jovanovic and Ressler, 2010; de Bitencourt et al., 2013). The effectiveness of these therapies underscores the dysfunction in controlling the unconscious fear response. Thus, Pavlovian fear conditioning is a valid preclinical tool used to explore the etiology of anxiety and fear-related disorders. In both humans and rodents, there are sex differences in cued and contextual fear conditioning as well as fear extinction learning. Males tend to exhibit higher levels of fear responses (i.e. freezing) than females; however this difference is influenced by ovarian estrogen levels (Colon and Poulos, 2020; Dalla and Shors, 2009; Day and Stevenson, 2020; Trask et al., 2020). Specifically, ovariectomized adult female rodents display the same level of fear as male rodents with estrogen replacement, resulting in lower freezing levels (Milad et al., 2009). Cycling estrogen also affects fear extinction as female rats in proestrus show enhanced extinction performance and memory compared to females in metestrus and males (Chang et al., 2009; Day and Stevenson, 2020; Milad et al., 2009; Perry et al., 2020). Thus, there is ample evidence that estrogen contributes to sex-specific fear behavior.

The endocannabinoid (eCB) system modulates behavioral phenomena including learning, memory, cognition, mood, stress and anxiety (TT-Y Lee et al., 2015; Lutz et al., 2015; McEwen et al., 2015; Moreira and Wotjak, 2010; Volkow et al., 2017). eCBs are particularly involved in the processing of context-dependent emotional valence (Lutz et al., 2015; Moreira and Wotjak, 2010; Volkow et al., 2017). Cannabinoid receptor 1 (CB1) is densely located in many brain areas associated with the limbic system, such as the hippocampus, cortex, amygdala, basal ganglia, nucleus accumbens and hypothalamus. Many of these structures are part of the HPA axis. The dense presence of CB1 in the limbic system and HPA axis implicates eCB signaling in the pathophysiology of stress-related mental disorders such as Major Depressive Disorder and PTSD (Lee et al., 2016; Lutz et al., 2015; McEwen et al., 2015; Moreira and Wotjak, 2010). The eCB system also significantly contributes to aversive learning, memory, and extinction. In male rodents, blocking CB1 enhances fear learning and memory recall, whereas exogenous CB1 activation reduces fear responses. Conversely, CB1 antagonism or genetic deletion impairs fear extinction while CB1 activation enhances this behavior (Moreira and Wotjak, 2010; Reich et al., 2008; Reich et al., 2013). Despite the overwhelming data in male rodents and humans, there is a paucity of data on the effects of eCB signaling on female fear behavior.

Sex differences in the eCB system are well established. In humans and animals, males tend to exhibit greater CB1 densities than females in most cortical areas (Gorzalka and Dang, 2012; Rubino and Parolaro, 2011); although female CB1 receptors are observed to have increased G-protein activation compared to male CB1 (Craft et al., 2013; Gorzalka and Dang, 2012; Rubino and Parolaro, 2011). Sex differences in CB1 levels are concomitant with sex-specific eCB modulation of hippocampal neurotransmission. Tonic and estrogen-driven anandamide (AEA) are present at female, but not male perisomatic and dendritic GABAergic-pyramidal cell synapses in rats (Huang and Woolley, 2012; Lee et al., 2010; S-H Lee et al., 2015; Tabatadze et al., 2015). Furthermore, at dendritic synapses, constitutive CB1 activity and tonic 2-Arachidonylglycerol (2-AG) modulates GABAergic neurotransmission in females and glutamatergic neurotransmission in males (Ferraro et al., 2020).

Exposure to chronic mild stress (CMS), a valid animal model of depression (Willner, 2005), reverses the sex differences in both hippocampal CB1 receptor levels and eCB signaling (Ferraro et al., 2020; Reich et al., 2013; Reich et al., 2009); suggesting that the eCB system is organized in male and female animals to respond differentially to chronic stress. In male rats, CMS enhances hippocampal-dependent trace-fear conditioning but does not affect hippocampal-dependent contextual fear conditioning. Exogenous CB1 activation decreases trace cued fear memory recall only in stress rats, whereas CB1 activation facilitates extinction in both stress and non-stress rats (Reich et al., 2013). Female rats were not included in this study, and it remains unclear if sex differences in CB1 levels and eCB signaling translate into behavioral phenomena. Therefore, the purpose of this study is to assess the effects of CMS and CB1 activation on hippocampal-dependent fear conditioning in female rats.

## Materials and Methods

### Subjects

Female Sprague-Dawley rats (Charles River, Boston, MA) were group-caged (3 per cage) and allowed to acclimate for 5-7 days prior to experimental testing. All animals were 40-43 days old at the beginning of experimental procedures and maintained on a 12-h/12-h light–dark cycle with lights on at 8:00 a.m. Food and water were available *ad libitum* in the home cages, unless otherwise noted. All experimental procedures were conducted in accordance with protocols established by the Institutional Animal Care and Use Committee of the Ramapo College of New Jersey.

### Chronic Mild Stress Protocol (CMS)

Animals were subjected to either the CMS protocol or the non-stress protocol (handled daily). Each day, 1-3 stressors were administered according to a set schedule. The complete regimen lasted 7 days/wk. for 3 wks. This protocol is modeled after Willner (2005) in that no individual stressor is considered severe, but the unpredictability of the protocol constitutes much of the stress. The stressors were: 1) 5 min swim in 20° C water, 2) cage rotation (social stress), 3) 18-hr social isolation with damp bedding, 4) 14-hr food deprivation, 14-hr water deprivation or 14-hr food and water deprivation, 5) 30 min physical restraint, 6) 30 min strobe light exposure and 7) 3-hr cage tilt. To minimize stress on the control animals, they were acclimated to social isolation prior to testing. We observe that this CMS protocol results in decreased body weight gain and reduces sucrose preference in both female and male animals (Reich et al., 2009, Reich et al, 2013). These effects are in accordance with the published behavioral effects of CMS protocols (Hill et al., 2005; Willner, 2005).

### Apparatus

Experiments were performed using an automated, computerized, fear-conditioning system (Coulbourn Instruments, White Hall, PA). The system consists of three conditioning chambers (30.48 x 25.4 x 30.48 cm) with removable stainless-steel grid floors. Footshocks are delivered through the floor grid via a shocker-scrambler unit controlled by custom-designed software (Coulbourn Instruments, White Hall, PA). Locomotor and freezing activities were monitored through CCD video cameras mounted at the top of each chamber and subsequently analyzed via FreezeFrame software (Coulbourn Instruments, White Hall, PA). A house light and speaker (1000 Hz tone, 80 dB) are located on the sidewalls of each chamber. In addition, each chamber is contained in a sound-attenuating cubicle equipped with a ventilation fan (60 dB).

### Trace Fear Conditioning

All fear conditioning procedures were performed in an isolated testing room. Animals were transported in their home cages to the testing room. The cages were covered to blind the animals during transport to and from the colony room.

*Acquisition:* For trace fear conditioning, each animal had a 120 sec baseline acclimation period before receiving three presentations of a tone CS (15 sec) followed by a footshock US (2 sec, 0.6 mA) that occurred 30 sec after the CS offset (i.e., trace period). CS-US presentations were separated by 180 sec inter-trial-intervals (ITI).

*Recall/Extinction:* The strength of the CS-US association was assessed 24 and 48 hours after the conditioning sessions. Trace retention/extinction trials consisted of a 60 sec baseline acclimation period followed by five trials of CS only presentation at 180 sec ITI. These spaced extinction trials were designed to minimize habituation to the altered (smaller, see below) chamber. During Recall Test #1 (24 hrs after Acquisition), *short-term extinction* was measured as the difference in freezing between Trial 1 and Trial 5. *Long-term extinction* was defined as the difference in freezing between Trial 1-Recall Test #2 (48 hrs after Acquisition) and Trial 1-Recall Test #1. For recall/extinction trials, inadvertent contextual conditioning was minimized by altering the chambers during testing sessions. The grid floor was replaced with smooth Plexiglas and another piece of Plexiglas was used to bisect the chambers (Manz et al., 2018; Reich et al., 2008; Reich et al., 2013). The outer walls were covered in multi-colored construction paper to further eliminate spatial cues. The chambers were cleaned with orange-scented 409 cleaner and the chamber assignments were shuffled from the previous day.

### Contextual Fear Conditioning

A conventional “foreground” contextual conditioning was used. The “foreground” protocol consisted of placing animals in the chambers for ten minutes and presenting three shock presentations (2 sec, 0.6 mA, 30 seconds apart) during the last two minutes. Immediately after the end of the session, each animal was returned to its home cage in the colony room. Each chamber was then carefully cleaned with Formula 409 cleaner at the end of each session. For recall/extinction tests, animals were placed in the same unaltered chamber used for conditioning. Once placed in the chamber, freezing was monitored for a ten-minute period. Initial fear memory recall (Recall Day 1) and subsequent long-term extinction recall (Recall Day 2) were assessed during the first two minutes of the ten-minute chamber access period. Short-term extinction was determined by comparing the last two minutes of the initial recall test (Time 5) with the first two minutes (Time 1).

### Pharmacological Treatment

Subgroups of stress and non-stress animals were either injected with the CB1 agonist ACEA, *N-(2-Chloroethyl)-5*Z*,8*Z*,11*Z*,14*Z*-eicosatetraenamide*, (0.1 mg/kg, i.p., Tocris Cookson, St. Louis, MO) or Vehicle (physiological saline and dimethylsulfoxide, 3:1 ratio) solution 30 min prior to experimental procedures. See Table 1. In previous studies, we tested the effects of three varying doses of ACEA (0.1 mg/kg, 0.5 mg/kg, and 1.0 mg/kg) on the locomotor activity of non-stress control animals. Activity was assessed by placing animals in the conditioning chambers for 15 min. Only the lowest dose of ACEA (0.1 mg/kg) did not significantly affect locomotor activity when compared to vehicle-treated animals; both groups showed ≤1 % immobility during the entire test period. A drug-induced increase in immobility may obfuscate changes in freezing behavior; therefore, the 0.1mg/kg dose was used in subsequent pharmacological experiments (Reich et al, 2013, Manz et al., 2018).

**Tabel 1:**
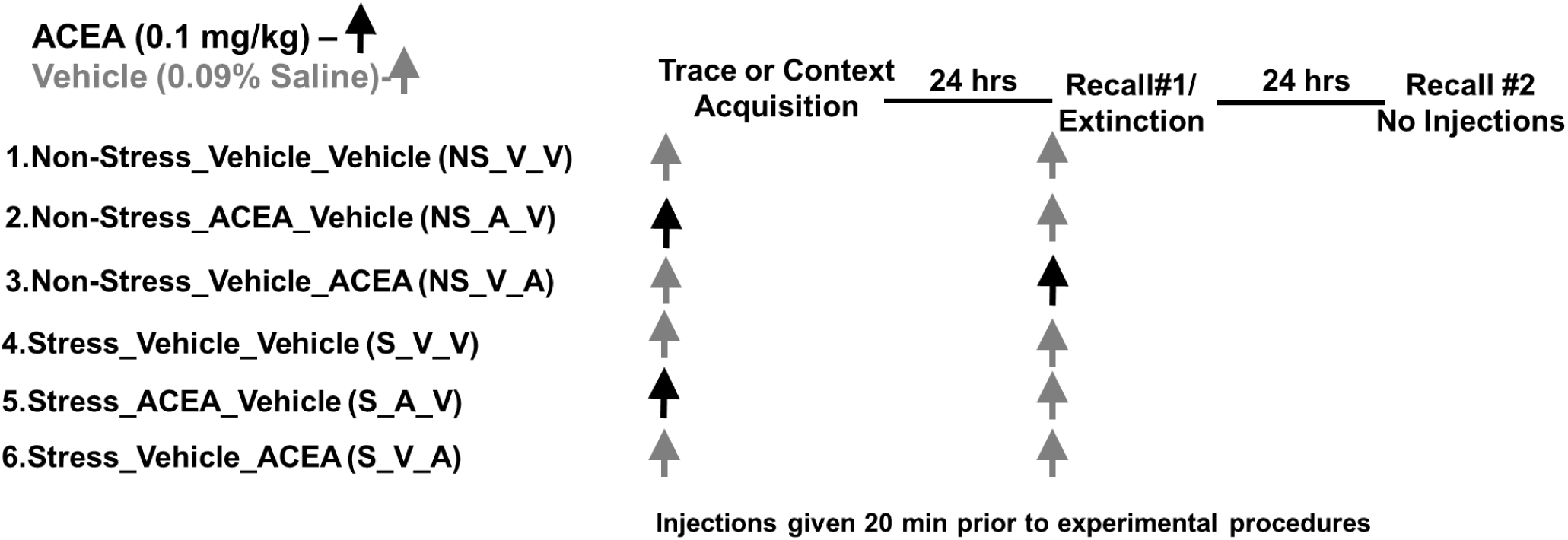
ACEA Experimental Protocol.

### Behavioral Measure and Data analysis

Freezing is defined as the absence of all movement except for respiration (Fanselow, 1980), lasting a period ≥ 3 sec. For all experimental sessions, time was binned into 15 sec intervals and the percentage of time spent freezing was calculated. If an animal had baseline freezing scores ≥ 2.5 standard deviations from the group mean it was removed from further analysis. In this study, no animals were removed from analysis. For the trace recall/extinction trials, freezing scores in 15 sec bins were averaged over the first minute following the CS offset. We observe that peak freezing to a CS tone occurs during this 60 second period (Reich et al., 2008, Reich et al., 2013, Manz et al., 2018). In trace fear recall tests, the first trial was used as an index of recall performance. Contextual recall was analyzed using average freezing during the first 120 sec of 10-minute chamber access time. Rescorla-Wagner learning theory suggests that these trial/time periods reflect the strongest potential for memory recall with later trial/time periods serving as extinction trials (Bouton, 2007). Data analysis (SPSS) was performed by repeated measures, one-way and two-way ANOVAs of group means with Tukey HSD post-hoc, pair-wise comparisons or Student-t-tests (p ≤ 0.05) as indicated.

## Results

### CMS increases both generalized and cued trace freezing behavior

Effects of CMS on fear acquisition and recall were assessed by a memory retention test performed 24 hrs after the conditioning. A two-way (Group x Trial) repeated measures ANOVA indicated that both stress (S) and non-stress (NS) female rats exhibited significant increases of cued freezing (60 seconds post 1^st^ CS) compared to baseline freezing (initial 60 seconds in chamber), (F (1, 6) = 45.95, p = 0.000, Fig 1a). CMS significantly increased cued freezing compared to NS controls (75.80 ± 3.42% vs. 58.20 ± 1.50%, F (1, 6) = 6.24, p = 0.047).

**Figure 1.**
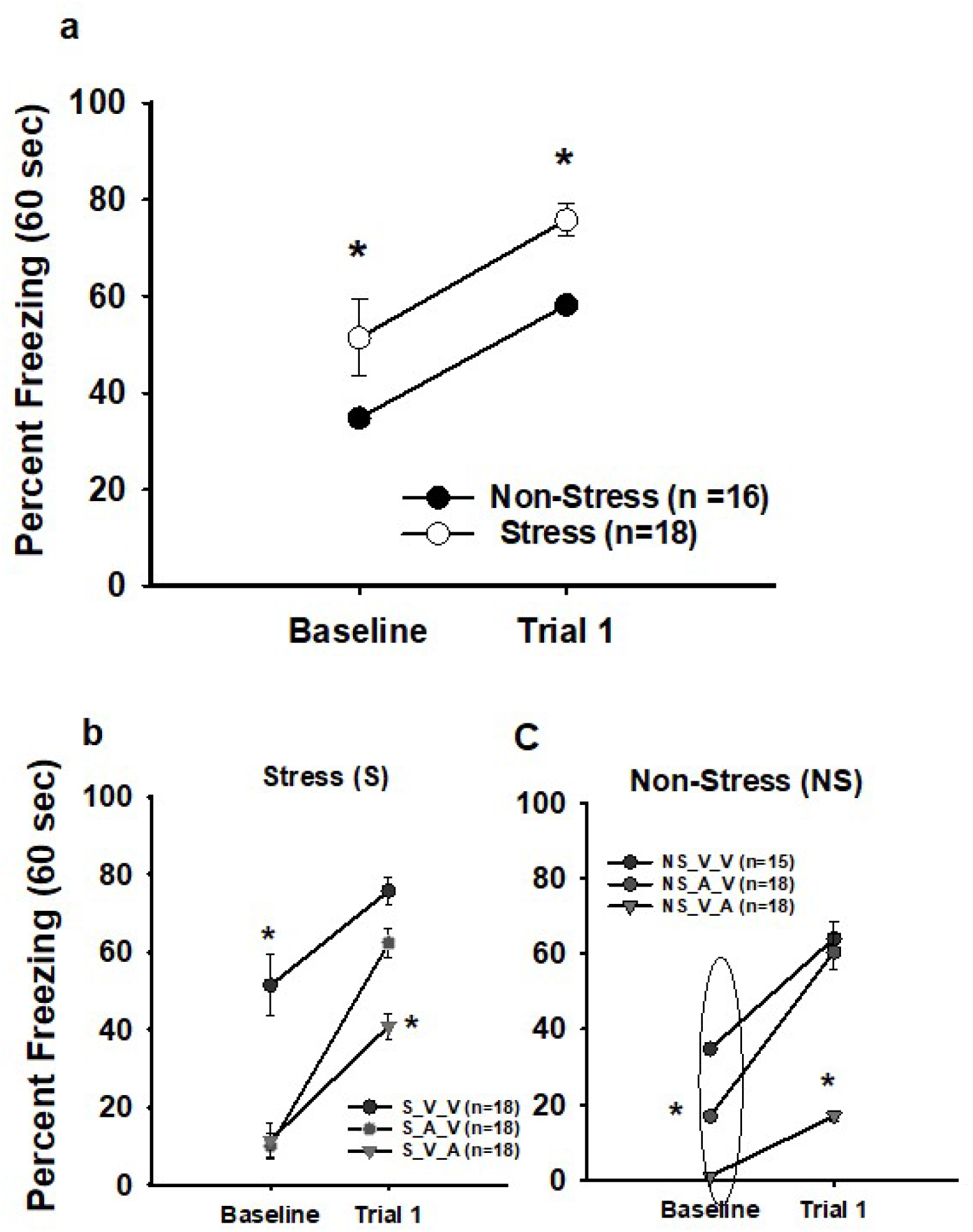
CMS enhances trace-fear conditioning in females. **a)** Baseline and cued freezing during the first trial of memory recall (60 sec post CS) are significantly greater in stress (S) vs. non-stress (NS) trace conditioned female rats. Both S and NS animals exhibit greater cued than baselined freezing. **b and c)** The CB1 agonist, ACEA, injected prior to both acquisition (S_A_V, NS_A_V) and recall (S_V_A, NS_V_A) reduces baseline freezing. However, only the S_V_A groups display less freezing during recall. Symbols represent mean ±SEM percent freezing. Asterisks indicates significant differences among groups during either the Baseline or Trial period (p ≤ 0.05).

Differences in baseline freezing rates also were observed between the two groups (S = 51.40 ± 7.98% vs. NS = 34.76 ± 1.51%, F (1, 6) = 6.24, p = 0.047). Baseline freezing is an index of a generalized response following aversive conditioning and may reflect the current anxiety level for an organism (Baldi et al., 2004; Jasnow et al., 2017). These observations indicate that CMS enhances both generalized and cued freezing behavior. An important note is that all rats received injections of physiological saline prior to conditioning and memory recall protocols. Since injection stress may affect fear conditioning and freezing behavior, these injections allow equitable comparisons to the subsequent experiments presented in this paper.

### CB1 receptor activation alters trace freezing behavior in both S and NS females

Prior to conditioning, S and NS rats were each split into three groups according to the protocol found in Table 1. In S females, a two-way (Group x Trial) repeated-measures ANOVA indicated a significant effect of Trial (F (1, 9) = 307.62, p = 0.000), indicating changes in freezing from baseline to Trial 1 across the groups. See Fig 1b. Differences also were observed during baseline freezing (F (1, 9) = 18.7, p = 0.001), indicating that ACEA modulates generalized freezing. A significant interaction effect of Group also was observed. Specifically, the S_V_V group (75.80 ± 3.42%) froze more during Trial 1, compared to the S_V_A group (40.90 ± 3.26%, p = 0.003, post-hoc analysis). No differences were observed between the S_V_V and S_A_V groups (62.34 ±3.82, p > 0.05).

For NS females, a two-way (Group x Trial) repeated-measures ANOVA revealed significant increases in freezing between baseline and Trial 1 (F (1, 9) = 105.95, p = 0.000; Fig 1c). A Group x Trial interaction and post-hoc analysis exposed significant differences in both baseline and cued freezing across the groups. During Trial 1, NS_V_V and NS_V_A (64.00 ± 4.41% vs. 16.94.00 ± 1.51%, respectively, p = 0.000) differed, although NS_A_V (60.44 ± 4.67%) and NS_V_V groups did not. ACEA also produced significant baseline freezing differences across all three groups.

### Effects of CB1 activation on trace-fear extinction

*Short-term extinction (STE):* During Recall Test #1, effects of CMS and ACEA on short term extinction were assessed by measuring the difference in freezing behavior between Trial 1 and Trial 5. For CMS exposed rats, a two-way repeated measures ANOVA (Group x Trial) resulted in a main Trial effect [F (1, 9) = 11.289, p = 0.008)] and a Group x Trial interaction effect [F (1, 19) = 19.98, p = 0.000)]. These effects indicate changes in freezing levels from Trial 1 to Trial 5 across the three groups. Post-hoc analysis revealed that only the S_V_V group showed significant STE from Trial 1 to Trial 5 (75.80 ± 3.42% vs. 46.88 ± 2.83%, p = 0.001). The S_A_V group displayed a nonsignificant decrease in freezing (62.34 ± 3.82% vs. 52.64 ± 3.52%, p > 0.05) and the S_V_A group increased freezing from Trial 1 to 5 (40.90 ± 3.26 %, vs 54.99 ± 1.61 %, p = 0.049). See Fig. 2a. In CMS females, ACEA administration either prior to trace fear acquisition or memory recall appears to impair STE relative to vehicle controls.

**Figure 2.**
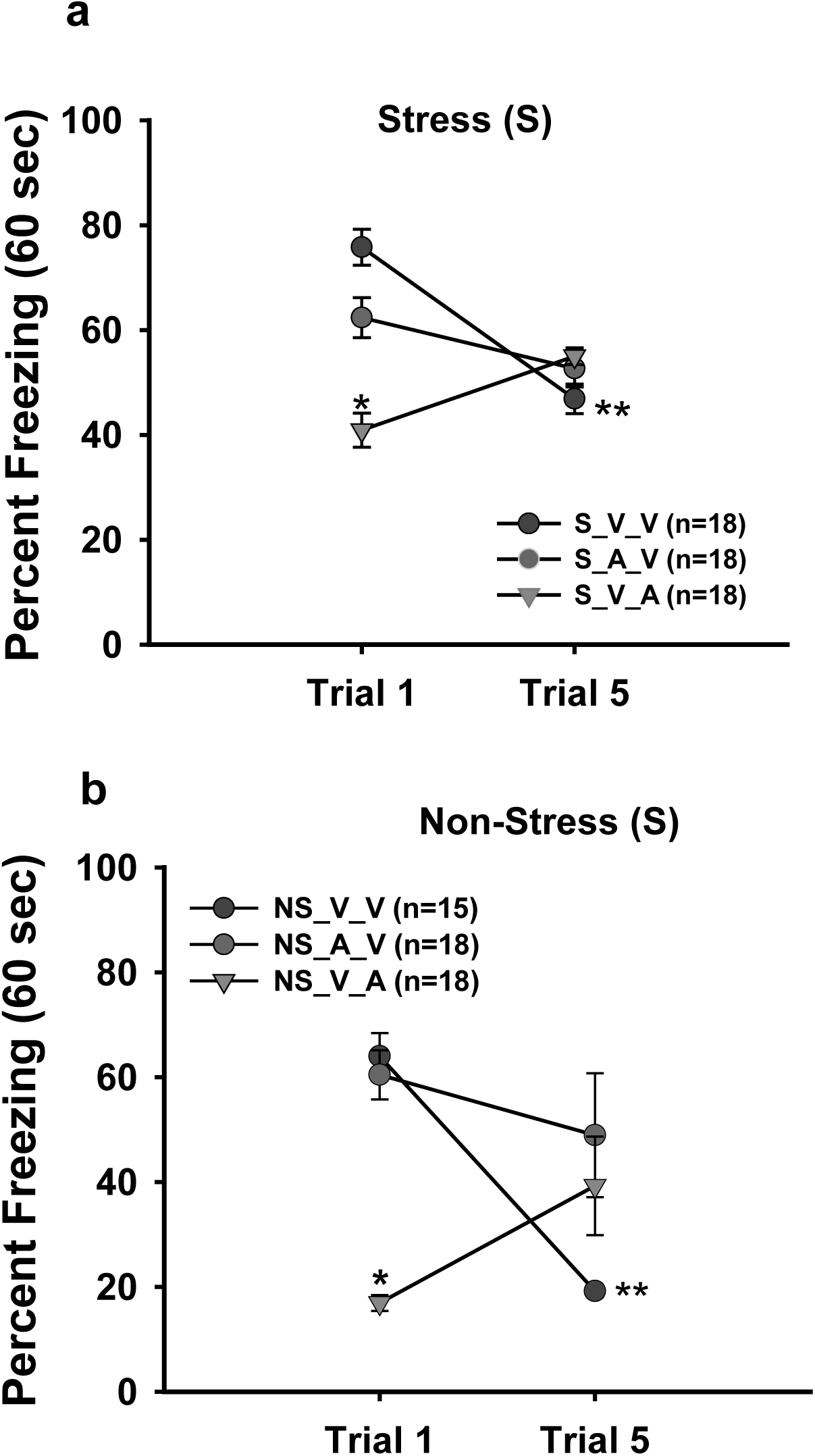
ACEA alters trace-fear short-term extinction (STE) in both S and NS females. **a)** and **b)**: In S and NS females, only the S_V_V and NS_V_V groups display significantly less freezing from Trial 1 to Trial 5. Administration of ACEA either prior to the Recall Test # 1 (S_V_A, NS_V_A) or prior to Acquisition (S_A_V, NS_A_V) impairs STE. Note that during Trial 5, the S_V_V and NS_V_V groups freeze less compared to either the S_A_V, NS_A_V, S_V_A or NS_V_A groups. Symbols represent mean ± SEM percent freezing. Single asterisk indicates differences among groups during Trial 1 (p ≤ 0.05). Double asterisks indicate significant changes in freezing from Trial 1 to Trial 5 per group.

In NS control females, a two-way repeated measures ANOVA (Group x Trial) indicates a Group x Trial interaction effect [F (1, 9) = 10.92, p = 0.007)], with no effect of Trial (p = 0.085). See Fig. 2b. Subsequent Post-hoc analysis showed only the NS_V_V group exhibited significant STE (64.00 ± 4.41% vs. 19.18 ± 1.41%, p = 0.003). Like the CMS condition, the NS_A_V group showed a small decrease in freezing (60.44 ± 4.67% vs. 48.92 ± 11.88%, p > 0.05) and the NS_V_A group increased freezing from Trial 1 to 5 (16.94.00 ± 1.51% vs. 39.26 ± 9.40%, p = 0.002). In both S and NS females, ACEA administration either prior to trace fear acquisition or memory recall appears to impair STE relative to vehicle controls.

*Long-term extinction (LTE):* To test if short-term extinction translated into long-term extinction, we assessed freezing 24 hrs following the initial recall testing (48 hrs following fear acquisition). For CMS females, A 2×3 (Recall Test x Group) repeated-measures ANOVA on first trial freezing resulted in a significant main effect of Recall Test (F (1,9) = 101.20, p = 0.000), indicating that there was a general reduction in freezing from Recall Test 1 (RT1) to Recall Test 2 (RT2) across the groups. Post-hoc analysis revealed that the S_V_V (75.80 ± 3.42% vs. 30.35 ± 2.88%, p = 0.001) and S_A_V (62.34 ± 3.82% vs. 37.45 ± 1.90%, p = 0.014) groups showed significant LTE, although group S_V_A (40.90 ± 3.26 %, vs 40.96 ± 3.30%, p > 0.05) did not display LTE (see Fig. 3a). A between-subjects ANOVA additionally resulted in differences during RT2 freezing (F (1,9) = 5.63, p = 0.026). The S_V_V animals froze significantly less than either S_V_V or S_A_V groups (p = 0.023). ACEA administration during initial recall/extinction appears to block LTE in CMS exposed female rats while during trace fear acquisition, ACEA moderately impairs LTE compared to stress vehicle controls.

**Figure 3.**
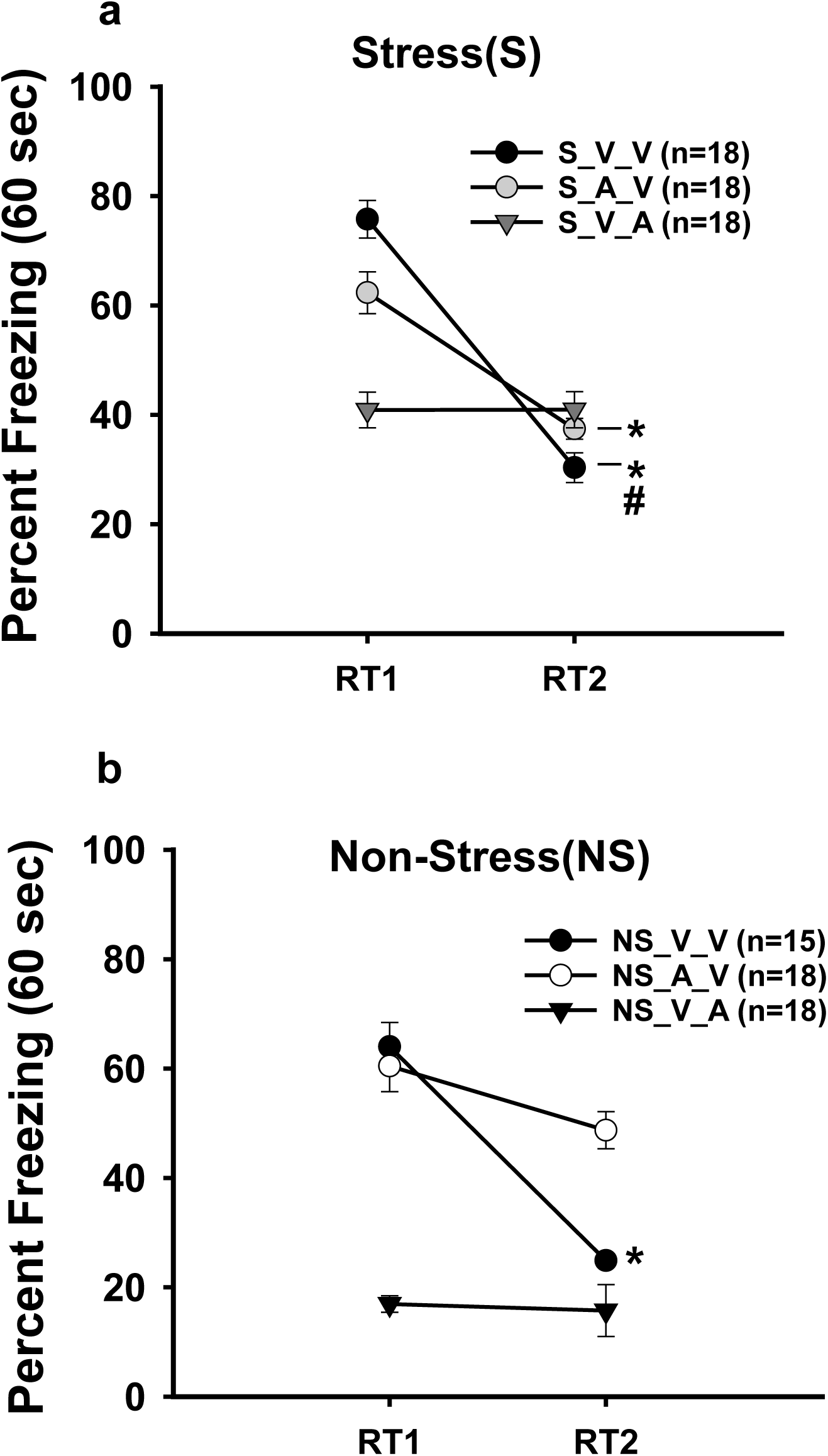
CB1 activation affects long-term, trace-fear extinction (LTE). **a)** S_V_V and S_A_V females show LTE, represented as significant decreases in freezing from Recall Test #1 (RT1) to Recall Test #2 (RT2). No changes in freezing occur for the S_V_A from RT1 to RT2. Asterisks indicate significant changes in freezing from Trial 1 to Trial 5 per group (p ≤ 0.05). The S_V_V froze the least during RT2 (pound sign), suggesting that ACEA (S_A_V and S_V_A) impairs LTE in CMS-exposed females. **b)** For NS females, only the NS_V_V group displayed significant LTE (asterisk, p ≤ 0.05), whereas neither the NS_A_V or NS_V_A freezing levels changed from RT1 to RT2. Symbols represent mean ±SEM percent freezing.

Analysis of NS controls with two-way 2×3 (Recall Test x Group) repeated-measures ANOVA showed a general reduction in freezing from RT1 to RT2 across groups Test (F (1,9) = 56.59, p = 0.000, Fig. 3b). Post-hoc analysis revealed significant decreases in freezing for both the NS_V_V (64.00 ± 4.41% vs. 24.88 ± 0.48%, p = 0.000) and NS_A_V freezing (60.44 ± 4.67% vs. 48.71 ± 3.39%, p = 0.033) groups. Like the CMS condition, the NS_V_A groups did not exhibit significant reduction in freezing from RT1 to RT2 (16.94 ± 1.51% vs. 15.73 ± 4.75%, p > 0.05). Between-groups analysis of RT2 indicated that all three groups displayed significantly different freezing levels (F (1,9) = 43.73, p = 0.000). The NS_A_V group froze the most, suggesting impairment of LTE.

### CMS does not alter contextual fear conditioning in females

In adolescent male rats, CMS exposure enhances hippocampal-dependent trace fear conditioning but does not affect contextual fear conditioning (CFC) (Reich et al., 2013). Experiments were performed to test if CMS would affect CFC in females. The strength of CFC was measured 24 hrs following acquisition by recording freezing during the first 120 sec of chamber access. No significant differences in CFC recall were detected between the NS and S groups (47.26 ± 4.24% vs. 41.46 ± 2.61%, p = 0.264, independent-samples t-test, Fig 4a). All rats received injections of physiological saline prior to conditioning and memory recall protocols.

**Figure 4.**
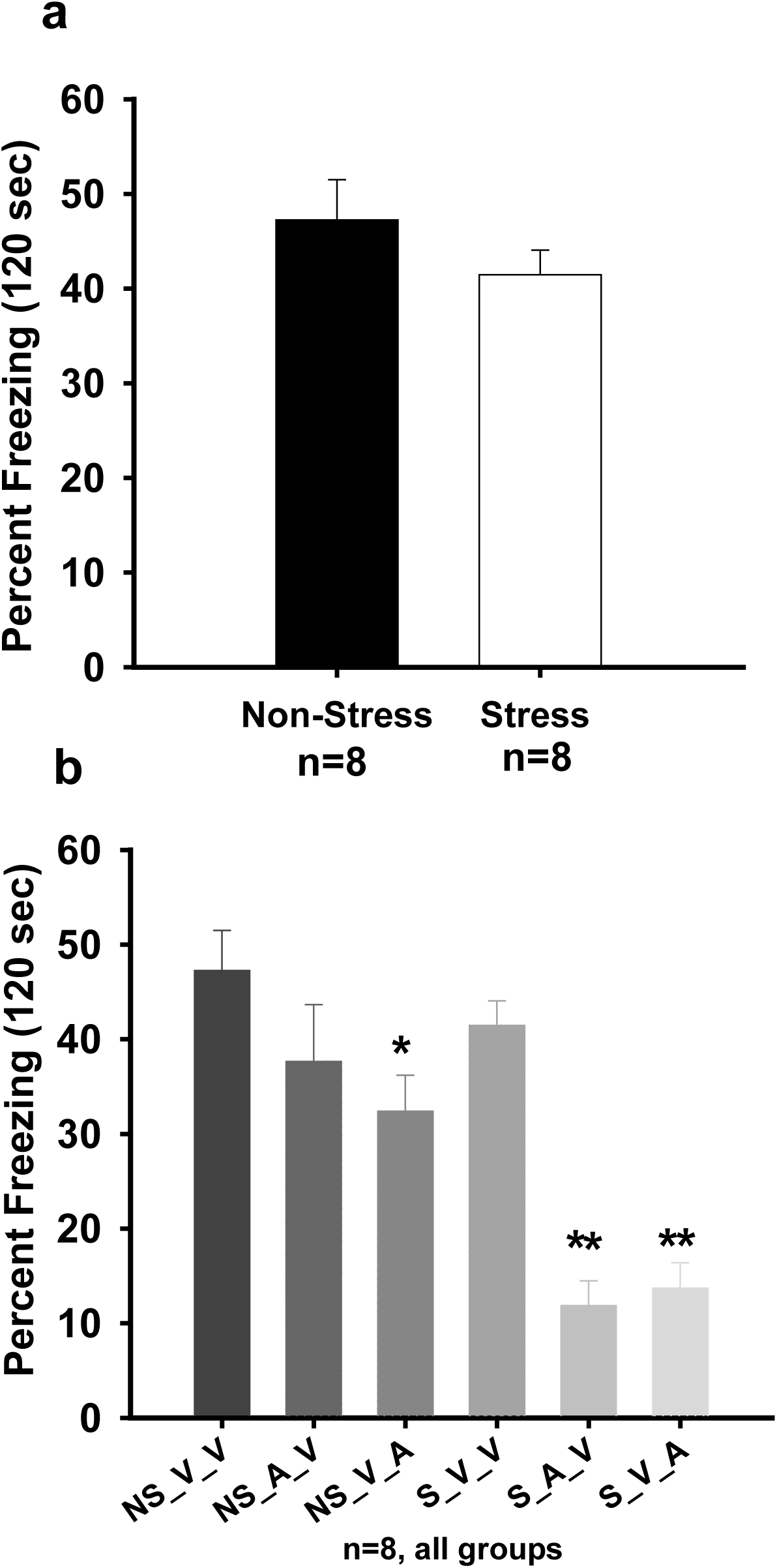
CB1 activation does affect contextual fear conditioning while CMS does not. **a)** Test of contextual memory strength 24 hrs following acquisition shows no difference between S and NS females**. b)** ACEA injection prior to recall decreases freezing for both the NS_V_A (single asterisk, p ≤ 0.05) and S_V_A groups (double asterisks, p ≤ 0.05). However, ACEA injection prior to acquisition decreases recall for the S_V_A group only. Freezing levels for both the S_A_V and S_V_A groups are significantly less than all other groups (double asterisks, p ≤ 0.05). Bars represent mean ±SEM percent freezing (120 sec).

### CB1 receptor activation differentially affects CFC in both S and NS females

ACEA (1.0 mg/kg) was injected according to the Table 1 protocol to assess effects of CB1 activation on CFC acquisition and memory recall. A one-way ANOVA indicated significant differences among S, NS and ACEA conditions (F (5,47) = 14.75, p = 0.000). Post-Hoc analyses confirm that NS_V_V and S_V_V controls do not significantly differ during recall. See Figure 4b. In the NS groups, the NS_V_A (32.40 ± 3.8%) females froze significantly less compared to either the NS_V_V (47.26 ± 4.24%) or NS_A_V (37.65 ± 5.95%) groups; however, NS_V_V and NS_A_V did not differ (p > 0.05). Comparatively, both the S_V_A (13.71 ± 2.69%) and S_A_V (11.88 ± 2.54%) groups froze significantly less than S_V_V (41.46 ± 2.61%, p = 0.000). S_A_V and S_V_A groups also differed significantly from all the NS groups (p = 0.001).

### Effects of CB1 activation on CFC STE and LTE

*Short-term extinction:* CFC STE is measured as the difference between the first 120 sec (Time 1 (T1) and the last 120 sec (Time 2 (T2)) of Day 1 recall/extinction. In the NS condition, a two-way repeated measured ANOVA (Group x Time) resulted in no main effect of Time (F (1,21) = 1.33, p > 0.05) and no main effect of Group (F(1,21) = 2.53, p > 0.05). Thus, no evidence of STE was observed for any of the NS groups. See Figure 5a.

**Figure 5.**
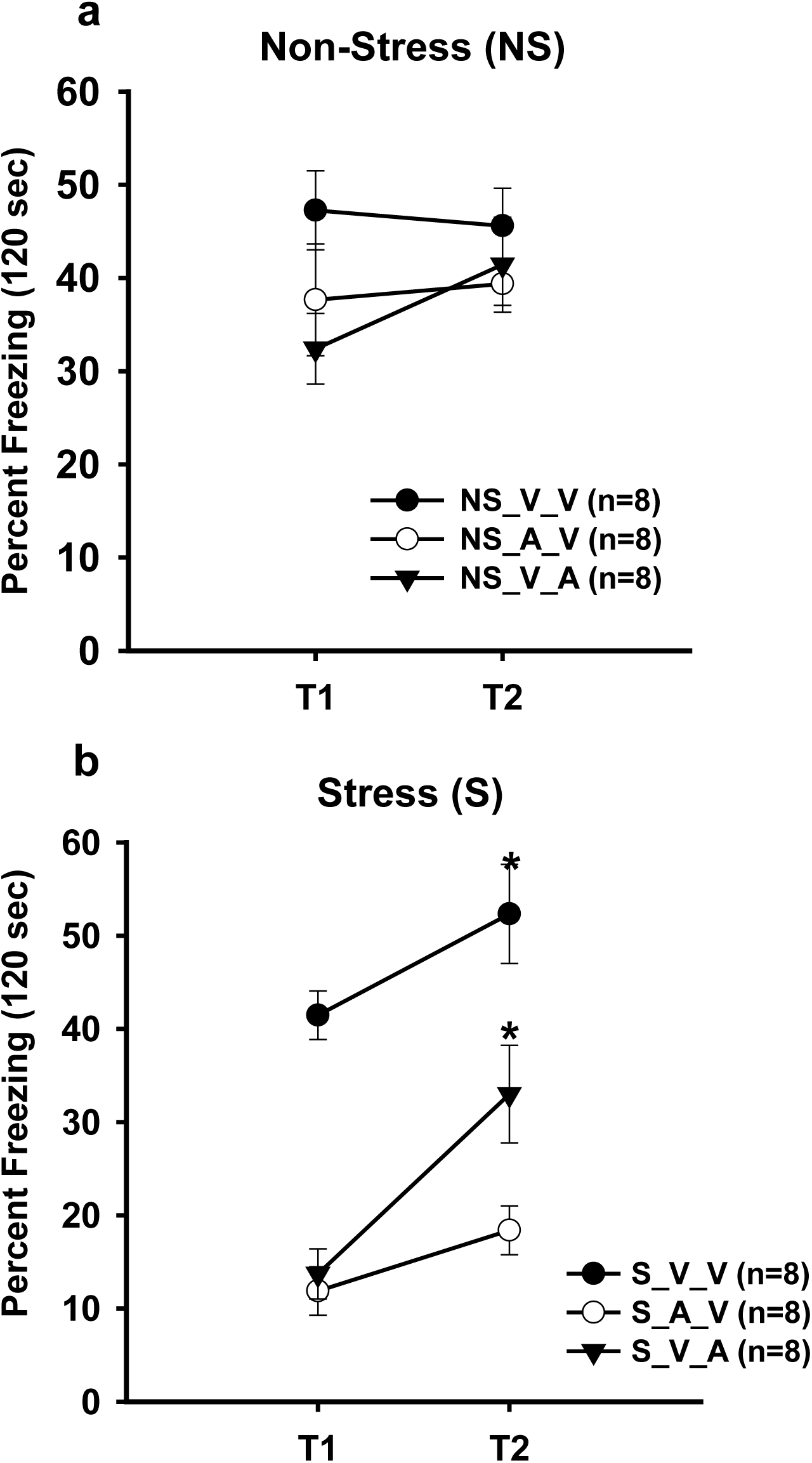
Impaired contextual short-term extinction in females. **a)** In the NS groups, freezing levels changed across the recall/extinction session (T1 vs T2). **b)** Both the S_V_V and S_A_V females displayed increases in freezing levels from T1 to T2 (asterisks, p ≤ 0.05). Freezing across the session did not change for the S_V_A group. Symbols represent mean ±SEM percent freezing.

For the S groups, a main effect of Time (F (1,21) = 27.64, p = 0.000) and a main effect of Group (F (1,21) = 50.58, p = 0.000 (two-way rmANOVA (Group x Time)) were observed. These findings indicate that significant differences between T1 and T2 were occurring across the three groups. See Figure 5b. Specifically, the S_V_V group froze more during T2 than T1 (52.33 ± 5.32% vs. 41.46 ± 2.61%, respectively: p = 0.05, pairwise contrast). The S_V_A group also froze more during T2 than T1 (33.09 ± 5.23% vs. 13.71 ± 7.60%, respectively: p = 0.002, pairwise contrast). The S_A_V did not display significant differences in freezing from T1 to T2 (11.87 ± 2.62% vs. 18.39 ± 2.62%, p > 0.05). These data show that STE did not occur for the S groups, although differences among the drug conditions were apparent for T2. The S_V_V maintained the highest levels of freezing while the S_A_V had the lowest levels. The S_V_A group displayed intermediate levels of freezing between S_V_V and S_A_V, suggesting that ACEA modulates conditioned freezing behavior across context exposure time in CMS exposed females. *Long-term extinction:* CFC LTE is measured as the difference between the first 120 sec during Day 1 recall/extinction (D1) and the first 120 sec during Day 2 recall/extinction (D2). A two-way repeated measured ANOVA (Group x Day) for the NS groups yielded a main effect of Day (F(1,21) = 16.974, p = 0.000) and main effect of Group (F(1,21) = 7.93, p = 0.003). See Figure 6a. Post-hoc analyses revealed that all groups displayed significant LTE (D1 vs D2): NS_V_V (47.26 ± 4.24% vs. 38.43 ± 2.58%), NS_V_A (32.40 ± 3.8% vs. 24.71± 4.24%) and NS_A_V (37.65 ± 5.95% vs. 21.56 ± 2.85%). The NS_V_A and NS_A_V groups froze significantly less during D2 than the NS_V_V group (p = 0.008); although the percent change of freezing from D1 to D2 was ∼8% for both the NS_V_V and NS_V_A groups. In contrast, the percent change for NS_A_V was 16%.

**Figure 6.**
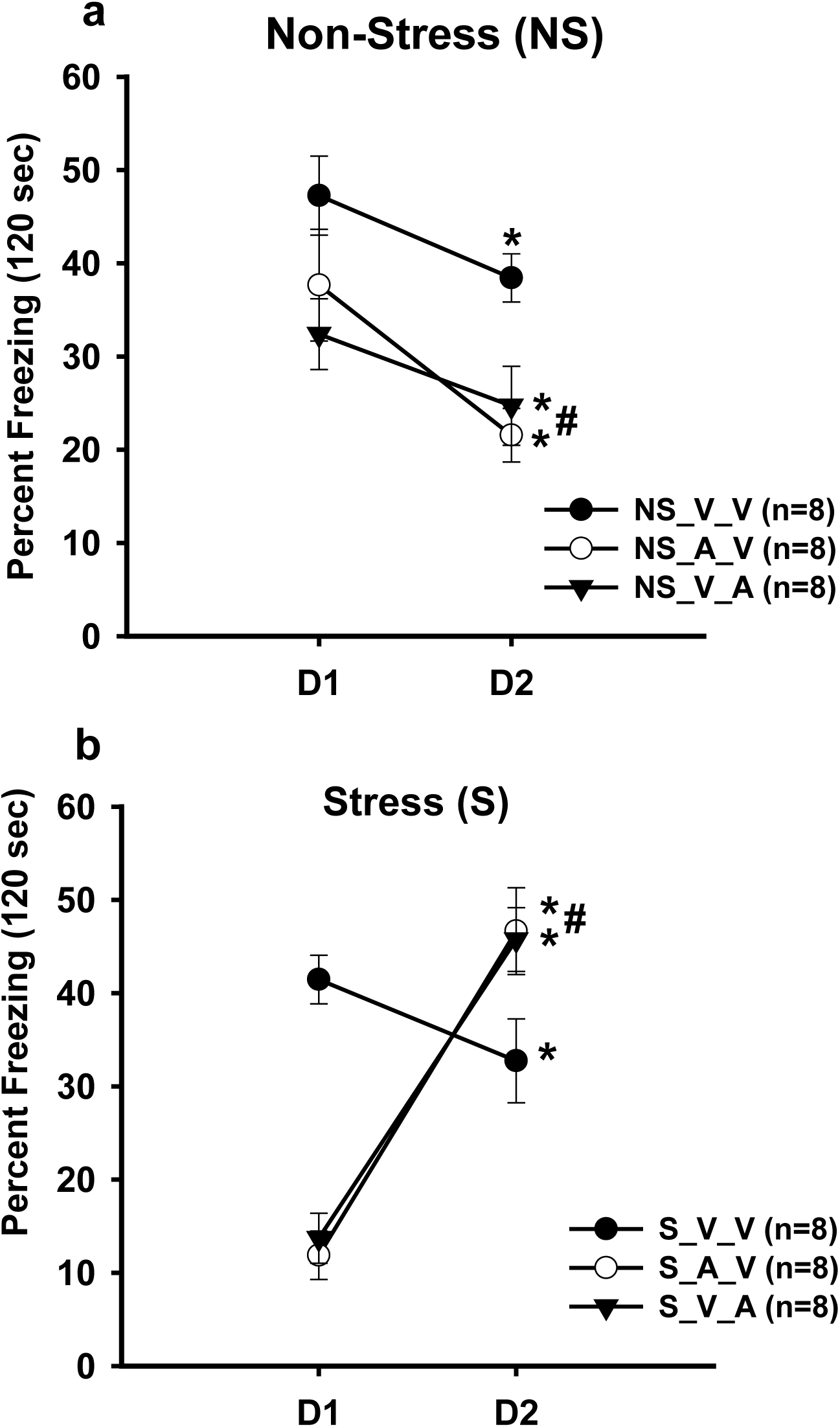
CB1 activation facilitates contextual long-term extinction in NS and impairs it in S females. **a)** All NS groups displayed significant freezing decreases from recall day 1 (D1) to recall day 2 (D2), asterisks (p ≤ 0.05). Both the NS_A_V and NS_V_A freeze significantly less than NS_V_V during D2 (pound sign, p ≤ 0.05). **b)** Only the S_V_V group exhibited significant LTE (froze less on D2 vs D1) with both the S_A_V and S_V_A groups showing increases in freezing from D1 to D2. Asterisks indicate differences from D1 to D2 per group (p ≤ 0.05). On D2, freezing for S_A_V and S_V_A groups are significantly greater compared to S_V_V (pound sign, p ≤ 0.05).

For the S groups, a two-way repeated measured ANOVA (Group x Day) resulted in a significant main effect of Day (F(1,21) = 39.75, p = 0.000) and interaction effect (F(1,21) = 20.64, p = 0.000). Specifically, the S_V_V froze less from D1 to D2 (41.46 ± 2.61% vs. 32.74 ± 4.5%), while both the S_V_A (13.71 ± 2.69% vs. 45.75 ± 3.43%) and S_A_V (11.88 ± 2.54% vs. 46.65 ± 4.65%) groups froze more. See Figure 6b. These observations suggest that in NS females, all groups exhibit LTE and ACEA administration prior to CFC acquisition may facilitate CFC LTE. However, in the S group, only the S_V_V group displayed LTE. ACEA administration in the S females appears to impair or reverse LTE.

## Discussion

The current data presents several key findings regarding the interaction of chronic stress and CB1 activation on hippocampal-dependent fear conditioning in adolescent female rats. Beginning with trace fear conditioning, CMS exposure enhanced freezing performance compared to NS controls, however CB1 activation modulates freezing behavior in both S and NS cohorts. ACEA administration prior to the first cued recall trial attenuated freezing similarly for S and NS females, whereas ACEA given prior to acquisition did not significantly affect cued recall. CB1 activation during initial trace fear memory recall may weaken the CS-US association in female rats, leading to less freezing behavior as observed in male rats (Lutz et al., 2015; Moreira and Wotjak, 2010; Reich et al., 2013). Alternatively, CB1 activation within the fear/defensive response circuitry could suppress freezing performance (Lutz et al., 2015; Moreira and Wotjak, 2010; Reich et al., 2008; Reich et al., 2013). Furthermore, CB1 modulation of baseline freezing in both S and NS groups suggests a reduction in *generalized* freezing behavior, which is associated with basal anxiety levels (e.g. higher basal anxiety = higher generalized freezing), (Baldi et al., 2004; Jasnow et al., 2017). Baseline freezing was attenuated by ACEA administration prior to both acquisition and recall, which indicates CB1/eCB recruitment lowers generalized anxiety during/following an aversive event in female rats. This finding contrasts with the effects of ACEA on trace baseline freezing in male rats (Reich et al., 2013); however, the CB1 antagonist/inverse agonist, AM251, increased trace baseline freezing in male mice (Reich et al., 2008).

For both S and NS groups, ACEA administration prior to acquisition or recall, either impaired trace STE or increased within-session freezing. This finding is consistent with the observation that 0.1 mg/kg ACEA impaired auditory-cued within-session extinction in intact, adult female rats (Simone et al., 2015). In contrast, CB1 activation in male adolescent rats did not affect trace STE and may have facilitated STE in CMS-exposed males vs. non-stress controls (Reich et al., 2013). Sex differences in eCB physiology may explain these notable comparisons between female and male trace STE. In the female hippocampal CA1, estrogen-mediated (ER_α_-mGLuR) AEA suppresses GABA release from perisomatic CB1 containing interneurons (Huang and Woolley, 2012; Tabatadze et al., 2015). Furthermore, females show both tonic AEA and 2-AG (Huang and Woolley, 2012; Tabatadze et al., 2015) activity at CA1 perisomatic synapses, whereas males show tonic 2-AG signaling only (Huang and Woolley, 2012; Lee et al., 2010; Lee et al., 2015; Tabatadze et al., 2015). We also recently reported evidence of an estrogenic-2-AG pathway and both tonic AEA and 2-AG in female CA1 dendrites (Ferraro et al., 2020). These physiological sex differences may contribute to the neural processes underlying either extinction learning or non-associative learning such as habituation and sensitization. eCB-mediated facilitation of extinction may depend on habituation-like phenomenon whereby, repeated presentations of a CS may allow activation of CB1 (Kamprath et al., 2006; Kamprath and Wotjak, 2004). Subsequently, repeated CB1 activation dampens signaling in the stimulus-response circuitry in male mice (Kamprath et al., 2006, 2011; Riebe et al., 2012). In adolescent females, sex-specific eCB physiology could lead to an opposite phenomenon where activation of CB1 results in CS sensitization. Buttressing this hypothesis, adolescent female Sprague-Dawley rats show impaired cued STE and LTE extinction during proestrus or met/diestrus (high estrogen levels) (Perry et al., 2020), whereas adult female rats display facilitated extinction during proestrus/ met/diestrus (Chang et al., 2009; Milad et al., 2009). Importantly, estrogen-dependent extinction requires estrogen receptor, Beta (ER_β_) (Chang et al., 2009). Perry et al. (2020) suggest that developmental differences in ER_β_/ ER_α_ densities contribute to shifts in adolescent and adult female fear extinction behavior. Although we did not track estrous cycles in the current study, we posit that ACEA administration alone or in combination with estrogenic ER_α_-eCB signaling promotes a sensitization process underlying the observed extinction impairments.

ACEA administration either prior to acquisition or recall, also impaired trace long-term extinction in both S and NS groups. In the NS females, CB1 activation prior to acquisition (NS_A_V) prevented robust decreases freezing as observed in the NS_V_V group. ACEA injections prior to initial memory recall (NS_V_A) did not yield any further freezing decreases from RT1 to RT2, although this simply may reflect a floor effect of freezing performance across the two testing sessions. For the S groups, the S_A_V group displayed significant LTE but was less than the S_V_V group. The S_V_A group did not exhibit LTE. Given the well documented role of CB1 and the eCB system in modulating extinction of aversive memories in male animals, these observations in females are surprising (Lutz et al., 2015; Moreira and Wotjak, 2010; C.G. Reich et al., 2008, 2013). These LTE impairments may reflect consolidated memories of the sensitized freezing observed during Recall Day 1’s extinction trials.

In male adolescent rats, CMS did not affect contextual fear conditioning (Reich et al., 2013). Our current observations also indicate that CMS exposure does not modulate contextual fear conditioning in adolescent female rats. Despite no evidence of a direct CMS effect on CFC, CB1 activation produced varied effects in both S and NS females. For NS rats, ACEA injection prior to the first recall/extinction session (NS_V_A) significantly attenuated freezing compared to rats treated with either vehicle (NS_V_V) or ACEA prior to acquisition (NS_A_V). In comparison, both the S_V_A and S_A_V groups froze less than the S_V_V group during the initial recall period. Furthermore, both the ACEA-treated S groups froze significantly less than all the NS groups. These observations suggest two mutual possibilities: 1) CMS does promote stronger CS_US association in CFC, even though freezing *performance* is initially equal for S and NS females and 2) CMS perturbations in female eCB/CB1 function discussed above (Ferraro et al., 2020, Reich et al., 2009) may underlie CMS strengthening of CFC. It is plausible that a CMS-induced hypofunction in hippocampal tonic eCB/CB1 signaling at GABAergic synapses biases synaptic gating towards stronger contextual-US associations. However, an increase in CB1 receptor density fostered by CMS may allow a stronger, exogenous CB1 activation to *rescue* eCB/CB1 function at hippocampal GABAergic synapses; thus, reducing the strength of CS_US relationship during acquisition or during memory recall. The observation that CFC in male rodents is impaired by exogenous activation of hippocampal CB1 receptors corroborates with this hypothesis (Arenos et al., 2006; Pamplona and Takahashi, 2006).

During the first CFC memory recall/extinction session, neither the NS and S groups exhibited significant STE, although end of session freezing levels were dependent on stress and drug exposure. All the S groups displayed increases in freezing from T1 to T2 with the S_V_A groups showing the largest increase. A similar within-session freezing increase was observed in the S_V_A and NS_V_A trace conditioned females, further suggesting that ACEA promotes non-associative sensitization during repeated CS only presentations. However, in CFC, CMS also appears to influence STE processes (either associative or non-associative).

Long-term CFC extinction was observed for all the NS groups. Notably, ACEA administration produced a modest gain in LTE for both the NS_V_A and NS_A_V groups compared to vehicle controls. For the S groups, only the S_V_V group showed LTE, while the ACEA groups displayed increases in freezing from RT1 to RT2. These freezing increases may reflect the consolidation of context-freezing memory at the end of the first recall/extinction session, where within-session freezing increased for the S_V_A and S_A_V groups. Regardless of mechanism, results from both the trace and contextual fear studies clearly indicate that compared to male rodents, CB1 activation during aversive extinction learning in females does not reliably facilitate extinction and appears to impair it under certain conditions.

ACEA activates the endovannilloid (eVN) transient receptor protein, TRPV1, as well as CB1 (Baker and McDougall, 2004; Casarotto et al., 2012; Fogaça et al., 2012), suggesting that our observed ACEA effects could arise from TRPV1 stimulation. Activation of brain TRPV1 increases fear and anxiety (Moreira et al., 2012). This is in stark contrast to CB1 activation; suggesting that the eCB and eVN systems regulate diametrically opposed actions on fear and anxiety at least in males (Moreira et al., 2012; Moreira and Wotjak, 2010). These behavioral differences are paralleled in male synaptic physiology; CB1 activation decreases glutamate neurotransmitter release, whereas TRPV1 activation increases it (Moreira et al., 2012). In the current study, if ACEA was acting though TRPV1 rather than CB1, then increases in freezing would be observed for both stress and non-stress rats during trace and contextual fear recall. Furthermore, the observation that ACEA decreased trace-conditioned baseline freezing favors a CB1 response over a TRPV1 response. A caveat is that in adolescent female rats, we observe CB1 activation increasing excitatory neurotransmission and TRPV1 activation decreasing it (Ferraro et al, 2020). Thus, the eCB and eVN systems may swap roles in females compared to males. Moreover, eVN could modulate anxiety (baseline freezing) and initial fear memory performance whereas the eCBs system regulates extinction-related processes. However, a recent study by Morena et al., (2020) suggests the opposite. In adult female rats, systemic AEA hydrolysis inhibition increased freezing behavior during cued fear extinction training and retrieval. The AEA effect was prevented by TRPV1, but not CB1 antagonism (Morena et al., 2020). Thus, in the current study, extinction impairment or enhanced freezing following ACEA administration may involve TRPV1 activation. In contrast to the current findings, Morena et al, did not observe eCB or eVn modulation of cued fear acquisition or initial memory recall. Age of rats (adolescent/young adult) vs. adult or experimental protocols (delay vs. trace and contextual conditioning) or pharmacological methodology could account for these differences. A tenable hypothesis is in that adolescent/young female rats, CB1 activation may modulate generalized and conditioned freezing behavior and TRPV1 activation may influence extinction learning and retrieval.

In conclusion, in adolescent female rats, we identify these main findings: 1) CMS enhances trace baseline and fear recall but does not affect contextual fear recall; 2) CB1 activation (ACEA) during either trace or contextual fear recall attenuates freezing behavior; 3) for trace conditioning, CB1 activation during Acquisition and Recall impairs both short and long term extinction regardless of stress exposure and 4) CB1 activation during contextual recall impairs both short and long term extinction in CMS exposed females only. Current and past observations from our lab clearly indicate that CMS (21-days) and CB1 activation affects trace fear recall in both male and female rats (Reich et al., 2013). However, in males, ACEA facilitates fear extinction but impairs it in females. These sex differences have implications in the use of medical cannabinoid treatment of stress-related anxiety disorders, such as PTSD as well as recreational cannabis use in adolescent/young adult females.

## Acknowledgements

We thank Gregory Mihalik, Cydney Mitchell, Amanda Swanson, Philip Sims, Timur Petrishin, Wendy Levine, Tayla Cacchione and Isabelle Weishaar, Zachary Mall, Brian Haner, Alexandra Kemp and Cecilia Wu for their amazing technical assistance. We also thank Michele Reich for proofreading.

## Funding Acknowledgements

This work was supported by National Institutes of Mental Health: grants RO3 MHO79294-01 and R15 MH085280-01 and Ramapo College Foundation Grants to Dr. Reich.

## Notes

### Competing Interest Statement

The authors have declared no competing interest.

